# Zika virus seroprevalence declines and neutralization antibodies wane in adults following outbreaks in French Polynesia and Fiji

**DOI:** 10.1101/578211

**Authors:** Alasdair D Henderson, Maite Aubry, Mike Kama, Jessica Vanhomwegen, Anita Teissier, Teheipuaura Mariteragi-Helle, Tuterarii Paoaafaite, Jean-Claude Manuguerra, W John Edmunds, Jimmy Whitworth, Conall H Watson, Colleen L Lau, Van-Mai Cao-Lormeau, Adam J Kucharski

## Abstract

**Background:** Serosurveys published following major outbreaks of Zika virus (ZIKV) have so far shown a high level of seroprevalence from samples collected within 12 months of the first confirmed case. A common assumption is that ZIKV infection confers long-term protection against reinfection, preventing ZIKV from re-emerging in previously affected areas for many years. However, the long-term immune response to ZIKV following an outbreak remains poorly documented.

**Methods:** We compared results from eight serological surveys, with sample sizes ranging from 49 to 700, before and after known ZIKV outbreaks in the Pacific region: five from cross-sectional studies of schoolchildren and the general population in French Polynesia over a seven-year period; and three from a longitudinal cohort in Fiji over a four-year period.

**Findings:** We found strong evidence of a decline in seroprevalence in both countries over a two-year period following first reported ZIKV transmission. In the cohort in Fiji, there was also a significant decline in antibody titres against ZIKV. However, the decline in seroprevalence was concentrated in adults, while high seroprevalence persisted in children.

**Interpretation:** The observed patterns of long-term anti-ZIKV antibody levels following outbreaks in the Pacific could be an early indication of the dynamics of population immunity in Latin America. Given that ZIKV antibody levels can wane substantially over time, follow-up seroprevalence studies and prospective clinical trial designs in Latin America may need to be revised, and assumptions about the potential for ZIKV to re-emerge may need to be revisited.

**Funding:** Pacific Funds, ANR, MRC, Wellcome, Royal Society.

## INTRODUCTION

Zika virus (ZIKV), a *Flavivirus* primarily transmitted to humans by *Aedes* mosquitoes, was first reported in the Pacific region on Yap island (Federated States of Micronesia) in 2007^1^. Six years later there was a large ZIKV outbreak in French Polynesia^2^ where an estimated 11·5% of the population visited healthcare facilities with clinical symptoms suggestive of ZIKV infection^3^. Since then the virus has spread across the Pacific region^4^, including to Fiji where cases of ZIKV infection were first detected in July 2015^5^. The same year, cases of ZIKV infection in Latin America were reported for the first time^6^. From February 1 to November 18, 2016, due to its rapid spread and association with birth defects, microcephaly in new-borns and Guillain-Barré syndrome in adults^7^ the WHO declared ZIKV a Public Health Emergency of International Concern^8^. At the end of 2016, outbreaks had declined in most of the countries recently affected^9^. However, ZIKV was still circulating in 2018 in several countries, including Fiji and Tonga in the Pacific region^10^.

In countries with known ZIKV outbreaks, the few serological surveys that have been published have demonstrated a high level of ZIKV seroprevalence following the outbreak. In French Polynesia, a population-representative cross-sectional serological survey at the end of the outbreak in 2014 suggested a seroprevalence of 49%^11^. In Martinique, a study of blood donors showed a post-outbreak seroprevalence of 42·2% in 2015^12^. In Salvador, Northeastern Brazil, a serosurvey in 2016 of prospectively sampled individuals including microcephaly and non-microcephaly pregnancies, HIV-infected patients, tuberculosis patients, and university staff, found a postoutbreak seroprevalence of 63·3%^13^. Finally, in paediatric and household cohort studies in Managua, Nicaragua, ZIKV seroprevalence was estimated to be 46% in households following the outbreak in 2016^14^.

It has been suggested that infection with ZIKV confers long-term immunity, which lasts several years. If so, the high level of seroprevalence in affected countries may provide herd immunity such that the current ZIKV epidemic is over in many locations and the virus will not be able to re-emerge for decades to come^2,9,13,15^. Recent evidence suggests that neutralizing antibodies can distinguish between ZIKV and dengue virus (DENV) – a closely related *Flavivirus* – and that the immune response following ZIKV infection can persist over a year^16,17^. Early data also suggests that primary ZIKV infection confers protective immunity^18^. However, ZIKV serosurveys conducted at the end of the outbreak and 18 months later in French Polynesia found a drop in seroprevalence in the Society Islands, the archipelago where over 85% of the inhabitants of French Polynesia reside^11^. Therefore, the long-term immune response following a ZIKV outbreak remains unclear. Consequently, so too does the potential for new outbreaks.

Here, we explore short- and long-term antibody responses against ZIKV as well as neutralizing response against ZIKV following two ZIKV outbreaks in the Pacific region. We compared results from five serological surveys in the Society Islands, French Polynesia, over a seven-year period, and three serial serological surveys in the same cohort of individuals in Central Division, Fiji, over a four-year period. These surveys span the pre- and post-outbreak period in each country, allowing us to examine temporal changes in antibody responses and hence, herd immunity, following a ZIKV outbreak.

## MATERIALS AND METHODS

### Study location and participants

#### French Polynesia

Four separate ZIKV serosurveys were previously conducted in the Society Islands (**Table 1**). As reported previously^11,19^, a first serosurvey (*n*=593) was conducted in adult blood donors recruited between July 2011 and October 2013, before the ZIKV outbreak that occurred between October 2013 and April 2014^3^. Two population-representative serosurveys were conducted among the general population, firstly between February and March 2014 (*n*=196), and then between September and November 2015 (*n*=700). An additional serosurvey was conducted among schoolchildren between May and June 2014 (*n*=476). Finally, a fifth serosurvey was conducted among schoolchildren in the Society Islands in June 2018 (*n*=457) using the same protocol as in 2014^11^.

**Table 1:** Seroprevalence of ZIKV among participants in five serological surveys in French Polynesia and three serological surveys in Fiji, conducted between July 2011 and June 2018

| Date | Country | Population | Age range (median) | Total no. seropositive/total no. tested | Seroprevalence % [95% CI] |
| --- | --- | --- | --- | --- | --- |
| Jul 2011-Oct 2013 | Society Islands, French Polynesia | Blood donors | 18-75 (36) | 5/593 | 0·8 [0·3-2]<sup>19</sup> |
| Feb-Mar 2014 | Society Islands, French Polynesia | General | 13-77 (47) | 18/49 | 37 [26-47]*<sup>11</sup> |
| Sep-Nov 2015 | Society Islands, French Polynesia | General | 4-88 (43) | 154/700 | 22 [16-28]*<sup>11</sup> |
| May-Jun 2014 | Society Islands, French Polynesia | School children | 6-16 (11) | 312/476 | 66 [60-71]*<sup>11</sup> |
| Jun-2018 | Society Islands, French Polynesia | School children | 6-16 (11) | 291/457 | 64 [58-69]*<sup>11</sup> |
| Oct-Nov 2013 | Central Division, Fiji | General | 2-78 (24) | 12/191 | 6·3 [3·3-11] |
| Nov-2015 | Central Division, Fiji | General | 4-80 (26) | 45/191 | 24 [18-30] |
| Jun-2017 | Central Division, Fiji | General | 6-82 (28) | 24/191 | 13 [8·2-18] |

\* CIs were calculated taking into account the cluster sampling design<sup>11</sup> and using the Fisher exact test.

#### Fiji

Two serial serosurveys were previously conducted in Fiji (**Table 1**). Individuals were first recruited into a population-representative community-based typhoid/leptospirosis seroprevalence study between September and November 2013^20^ (*n*=1,787), before autochthonous transmission of ZIKV was first detected in July 2015^5^. Briefly, nursing zones were randomly selected, from which one individual from 25 households in a randomly selected community was recruited. Participants who had consented to being contacted again for health research were subsequently recruited in November 2015 in 23 communities in Central Division through last known addresses, phone numbers and the assistance of local nurses (*n*=327). A third follow-up serosurvey was conducted in June 2017 using the same protocol as in 2015^21^ (*n*=321). Follow-up surveys were only performed in Central Division, which was the focus of a DENV outbreak in 2013/14^21^. Only blood samples serially collected from the same participants (*n*=191) in 2013, 2015 and 2017 were analyzed in the present study.

### Informed consent and ethics approvals

#### French Polynesia

The five serosurveys were approved by the Ethics Committee of French Polynesia (ref 61/CEPF 08/29/2013, 60/CEPF 06/27/2013, 74/CEPF 05/04/2018, and 75/CEPF 05/04/2018).

#### Fiji

The original 2013 study, and the 2015 and 2017 follow up studies were approved by the Fiji National Research Ethics Review Committee (ref 2013–03, 2015.111.C.D, 2017.20.MC) and the London School of Hygiene and Tropical Medicine Observational Research Ethics Committee (ref 6344, 10207, 12037).

### Serological analysis

#### French Polynesia

Serum samples collected from blood donors between July 2011 and October 2013 and samples collected from the general population and schoolchildren in 2014 were all tested for presence of IgG antibodies against ZIKV and each of the four DENV serotypes using a recombinant antigen-based indirect ELISA as reported previously^11,19^. Samples collected from the general population in 2015 and from schoolchildren in 2018 were tested by microsphere immunoassay (MIA) using the same recombinant antigens as for the ELISA^7,11^.

#### Fiji

All serum samples collected in Fiji were tested using MIA to detect IgG antibodies against ZIKV and all four DENV serotypes as previously reported^7,11,21^.

As an additional validation, a subset of samples collected in 2013 and 2015, and all samples collected in 2017 were tested for the presence of neutralizing antibodies against ZIKV and each of the four DENV serotypes using a neutralization assay as previously described^7^. Among the 14 individuals that tested seronegative against all viruses in 2013 and were re-tested in 2015, there was no evidence of an association between changes in ZIKV titre and changes in any of the DENV titre, suggesting that the changes in ZIKV titre were unlikely to be influenced by DENV cross-reaction (**Supp. Fig 1**).

### Statistical analysis

For data from Fiji, where serial samples were collected from the same individual, changes in seroprevalence between studies were tested using McNemar’s test. In French Polynesia, chi-squared tests were performed to test for evidence of a change in seroprevalence between two cross-sectional surveys. Changes in mean titre level between groups were analyzed using a t-test.

All data and code used in the analysis are available at: *https://github.com/a-henderson91/zika-sero-pacific.git*.

## RESULTS

ZIKV seroprevalence estimates from blood samples collected before and after the first confirmed autochthonous transmission of ZIKV in French Polynesia and Fiji are shown in **Table 1**. In French Polynesia, seroprevalence of anti-ZIKV IgG in blood donors recruited before October 2013 was <1%, which confirmed that the virus had not previously circulated in the population. The analysis of population representative samples collected in the Society Islands of French Polynesia after the emergence of ZIKV showed a decrease in ZIKV seroprevalence from 37% to 22% between February-March 2014 and September-November 2015 (chi-squared test, *p*=0·03). In Fiji, analysis of the serum samples serially collected from a cohort of participants in the Central Division showed an increase in ZIKV seroprevalence from 6·3% in October-November 2013 to 24% in November 2015 (chi-squared test, *p*<0·0001), and then a decrease to 13% by June 2017 (chi-squared test, *p*=0·008). Moreover, in this cohort, seven of the 191 participants seroconverted (from negative to positive) and 28 seroreverted (from positive to negative) to ZIKV between 2015 and 2017 (McNemar’s test, *p*=0·0007) (**Supp. Table 1**)

To investigate possible factors influencing the decline in seroprevalence, we compared the seroprevalence profiles in children (≤16 years) and adults (≥17 years) in both settings (**Table 1 and Figure 1**). In French Polynesia, although ZIKV seroprevalence declined in the general population from the Society Islands over 18 months, there was no evidence of a decline in seroprevalence in two serosurveys conducted four years apart in schoolchildren aged 6 to 16 years (chi-squared test, *p*=0·58) (**Table 1 and Figure 1**). In Fiji, in adults aged over 16 years, there was a decrease in seroprevalence from 24% in 2015 to 13% 2017 (**Figure 1**). There were two seroconversions in the collected samples over this period but 24 seroreversions (McNemar’s test, *p*<0.0001) (**Supp. Table 1**). In contrast seroprevalence in children remained stable over the period at 19% (**Figure 1**), with five seroconversions and four seroreversions (McNemar’s test, *p*=1) (**Supp. Table 1**).

**Figure 1:**
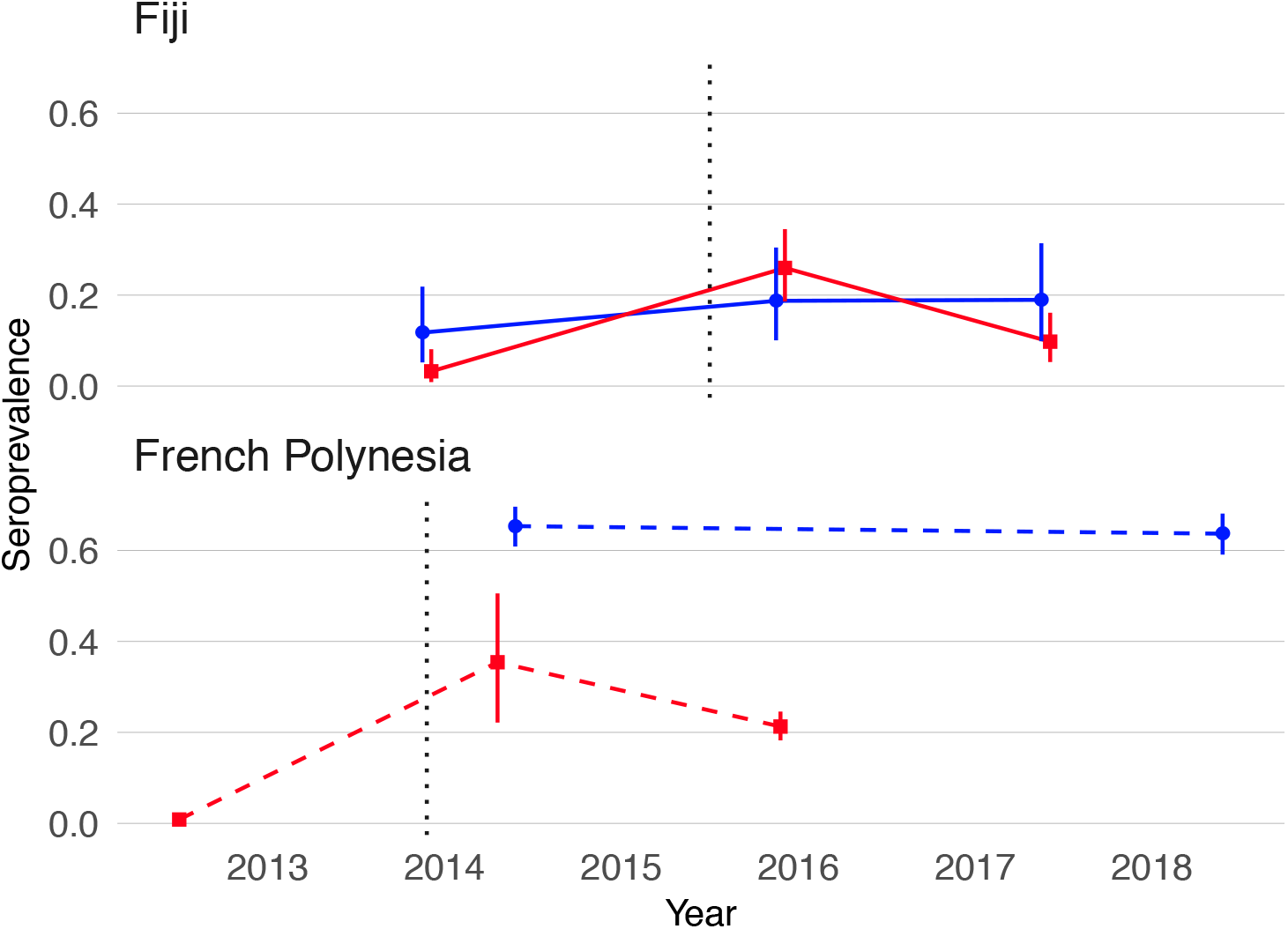
Decline in ZIKV seroprevalence following outbreaks in Fiji and French Polynesia, by age group. Blue, seroprevalence and 95% confidence intervals for children (aged ≤16 years). Red, seroprevalence and 95% confidence intervals for adults (aged ≥17 years). Solid lines, trends in data collected from the same individuals. Dashed lines, trends in seroprevalence between population representative cross-sectional surveys. Grey dotted lines mark the first confirmed ZIKV case in each country.

In order to assess whether the decline in ZIKV seroprevalence was also observed for other circulating *Flaviviruses*, the seroprevalence pattern against each of the four DENV serotypes was analyzed in both countries, by age group (**Supp. Figure 2**). In Fiji, seroprevalence for DENV-1, DENV-2 and DENV-4 increased in participants aged both under and over 16 between 2013 and 2017 (**Supp. Figure 2A, B and D**). DENV-3 seroprevalence also increased in both age groups between 2013 and 2015 following an outbreak in 2013-14^21^ and then declined in 2017 but, unlike ZIKV, this drop was not significant in participants aged under or over 16 years (McNemar’s test, *p*=1 and *p*=0·1213, respectively) (**Supp. Figure 2C**). In French Polynesia between 2014 and 2018, the seroprevalence in children aged under 16 years showed no evidence of a change for DENV-1 and DENV-2 (chi-squared test, *p*=0·1917 and *p*=1, respectively) and decreased for DENV-3 and DENV-4 (chi-squared test, *p*<0·0001 and *p*=0·0085, respectively). In participants from the general population, seroprevalence for all four DENV serotypes declined between 2014 and 2015 (**Supp. Fig 2A-D**). Despite similar age distributions in participants between the two surveys conducted in the general population in 2014 and 2015, a higher proportion of the samples in 2014 tested positive for all four DENV serotypes, suggesting that the sampling included a higher risk group for arbovirus infection than those sampled in 2015 (**Supp. Table 2**). To adjust for this potential bias, seroprevalence for the four DENV serotypes was estimated from a bootstrap sample of the 2014 responses, with replacement, weighted by the DENV exposure profile in the 2015 survey so that the bootstrap sample of the 2014 responses had a similar DENV exposure profile as in the 2015 responses (**Supp. Table 3**). After adjusting for this sampling bias, there was no evidence of a change in seroprevalence for any of the four DENV whereas the decline in ZIKV was still present (**Supp. Table 3**).

To explore dynamics of antibody waning at the individual level, we performed neutralization assays on a subset of 46 participants from Fiji for whom sufficient sera were available from all three collection periods. We found that in the 32 individuals who were ZIKV seropositive in 2015, anti-ZIKV antibody levels waned significantly in 2017, with an average decline in log titre of −1·94 (t-test,*p*<0·001) (**Figure 2A and Supp. Table 4**).

**Figure 2:**
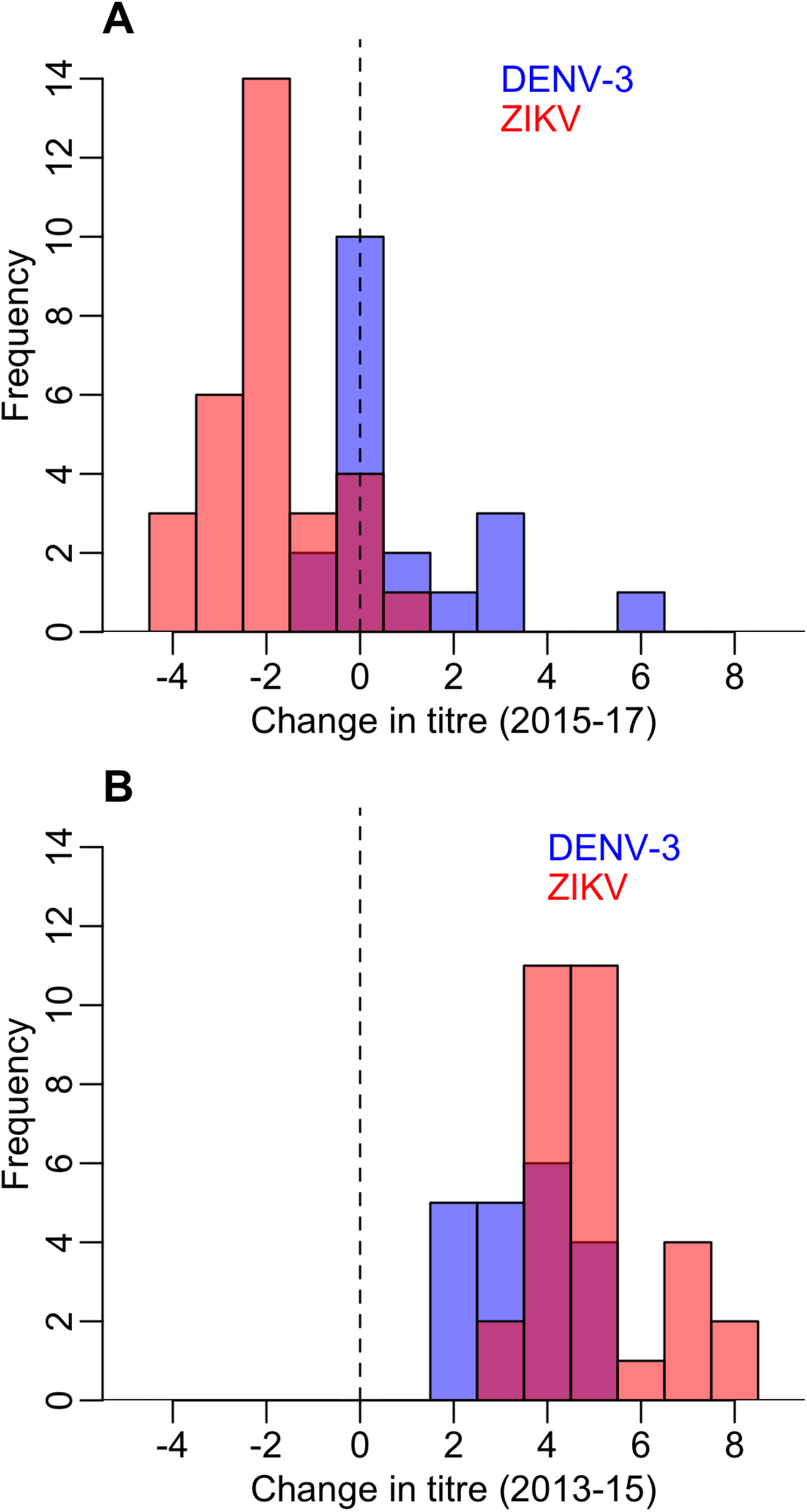
Waning of neutralizing antibody responses against ZIKV. A). Histogram of change in log titre against DENV-3 and ZIKV between 2015–2017 for individuals who seroconverted to these respective viruses between 2013–2015. B) Histogram of change in log titre against DENV-3 and ZIKV for individuals who seroconverted to these respective viruses during 2013–2015

To test whether the dynamics of anti-ZIKV antibody waning were different from the response to DENV infection, we compared results for ZIKV to the neutralization response following a DENV-3 infection in the same cohort from Fiji. There was a large DENV-3 epidemic between 2013-14 in Fiji^21^, which meant most seroconversions to DENV-3 occurred between the collection of samples in 2013 and 2015. In those individuals that seroconverted to DENV or ZIKV between 2013 and 2015, the initial rise in antibody log titres against ZIKV was larger than for DENV-3, with a change of 5 0 and 3·37 respectively (**Figure 2B and Supp. Table 4**).

However, while the mean antibody log titre for ZIKV declined between 2015 and 2017 (−1·94) (t-test, *p*<0·001), it increased for DENV-3 (089) over the same period (t-test, *p*=0·04) (**Figure 2A and Supp. Table 4**).

## DISCUSSION

Analyzing data from serological surveys conducted in French Polynesia and Fiji at different time points after the first reported autochthonous ZIKV transmission, we found strong evidence of a decline in ZIKV seroprevalence. The high number of participants from the Fijian cohort that seroreverted between 2015 and 2017 suggested that anti-ZIKV antibody levels waned in these individuals to the point that they were no longer detectable. Using a neutralization assay to test paired sera collected in Fiji, we found that the mean change in neutralizing antibody titres against ZIKV also decreased significantly between 2015 and 2017, showing that individual-level antibody titres against ZIKV as well as overall seroprevalence decreased over time. In contrast over the same period, neutralizing antibody titres against DENV-3, a closely related *Flavivirus* which caused a large epidemic in Fiji in 2013-2014^21^, remained stable.

In both countries we found seroprevalence against ZIKV in adults (aged over 16) declined over the two-year period following an outbreak, while the overall level of seroprevalence persisted in children. This pattern was unique to ZIKV compared to DENV in both countries. It is possible that this is related to the DENV immunological profile of individuals, given that the older population are likely to have experienced more DENV infections over their lifetime. If an individual has experienced prior DENV infections, high numbers of weakly neutralizing cross-reactive B cells may outcompete naïve B cells for ZIKV antigen^22^, leading to a short-term boost in antibody response against ZIKV^23^ but not a persistent specific response; a similar phenomenon has been observed for other antigenically variable viruses like influenza^24^. If this is the case, we might expect ZIKV antibody responses to wane more in individuals with prior DENV exposures. This trend can be observed in longitudinal data collected in Fiji where the relative decrease in neutralization titre between 2015 and 2017 was greater in individuals who tested positive for at least one DENV serotype (**Supp. Figure 3**), although the pattern is not conclusive since there is a lack of power to analyze this relationship due to the small number of available samples tested. To our knowledge, the only other study that investigated the longterm persistence of neutralizing antibodies against ZIKV was conducted in 62 residents of Miami (Florida, USA), who had a confirmed ZIKV infection in 2016^17^. This cross-sectional study found that all participants had neutralizing antibodies against ZIKV 12 to 19 months after infection. This study also found that at least 37% of the participants had no evidence of past DENV infection, which supports the hypothesis that anti-ZIKV immune responses may persist longer in populations that have had less exposure to DENV. More data are therefore needed to test the hypothesis about the potential impact of pre-existing anti-DENV immune response on anti-ZIKV antibody waning.

Although we found evidence of waning antibodies following two ZIKV outbreaks, the implications for susceptibility to further ZIKV infection remain unclear. Given the antigenic similarity of DENV and ZIKV^25^, it is commonly assumed that the immune response to ZIKV infection will be similar to that following DENV infection. High levels of neutralizing antibodies to DENV have been shown to correlate with protection from symptomatic infection^26^. Moreover, infection with a single DENV serotype can confer lifelong immunity to the infecting serotype as well as a transient period of cross-neutralization against heterologous serotypes^27^. However, it is unclear in the context of DENV, as well as ZIKV, what the relationship is between a specific titre value and susceptibility to further infection. Despite these caveats, our results show that anti-ZIKV antibody levels can wane substantially over time and long-term antibody dynamics following a ZIKV outbreak are distinctive to those following a DENV outbreak in the same location during the same period.

There are some additional limitations to our analysis. First, we did not have polymerase chain reaction (PCR) confirmation of ZIKV infection in individuals sampled in this study. We have presented analysis of representative serological surveys in two locations with known, PCR-confirmed ZIKV outbreaks^5,28^. However, PCR confirmation for ZIKV at the individual level remains difficult to obtain, in particular from blood samples, and there have been relatively few confirmations globally^28^, let alone analysis of long-term antibody dynamics in PCR confirmed patients.

Our analysis was also limited by study design. In French Polynesia, surveys were cross-sectional, so we were unable to examine temporal antibody dynamics at the individual level. However, both cross-sectional studies of the general population were conducted using population representative cluster sampling^11^ in the same remote island locations with stable population, which enabled robust comparisons of overall seroprevalence. We did identify one potential source of sampling bias with different DENV exposure profiles in the two surveys, but our conclusions of declining seroprevalence for ZIKV persisted once we adjusted for this bias (**Supp. Fig 3, Tables 1-2**). We also used different serological analyses between the studies in French Polynesia in 2014 and 2015, however both used the same recombinant antigens and it has been shown that there was good agreement between ELISA and MIA in the 2014 samples^11^. In Fiji, a strength of our study was the collection of longitudinal samples from the same individuals at three time points. However, our sample size was limited given the logistical challenge of recontacting participants twice over a four-year period. These data provided very strong evidence that ZIKV seroprevalence declined over the two-year period following first reports of circulation, but our sample size was insufficient to fully explore the potential effect of anti-DENV pre-existing immunity on anti-ZIKV antibody waning once we stratified individuals by previous DENV exposure.

The global ZIKV epidemic began in the Pacific islands in 2013 before spreading in Central and South America from 2015. Seroprevalence studies following ZIKV epidemics in Latin America have been reported but data have either been non-representative^13^ or not enough time had elapsed since the outbreak to observe longterm dynamics^14^. To our knowledge, these are the first studies of seroprevalence over a long-term period following a ZIKV outbreak. Therefore, patterns observed in Pacific islands may be an early indication of what might happen in Latin America where ZIKV outbreaks began two to three years after the French Polynesia epidemic^3,29^.

In the short-term, our findings have implications for the design of follow up studies of ZIKV in Latin America. Specifically, our results provide evidence that levels of seroprevalence one to two years following ZIKV circulation may be lower than previously expected and study designs may need to be adapted to reflect this. For example, estimates of microcephaly risk may be inflated if derived from long-term seroprevalence data that underestimate the true extent of infection within the population. There may also be a stronger case for pursuit of vaccine development if low seroprevalence and waning antibody levels against ZIKV are associated with susceptibility to reinfection, and hence future outbreaks. However, designs of vaccine trials^30^ could be affected if post-outbreak seroprevalence levels are lower than originally expected. Potential for waning of long-term responses would also need to be considered during vaccine development, as would the DENV exposure history of vaccine recipients. Finally, our results could have implications for public health planning in countries that have experienced ZIKV epidemics. Predictive modelling has typically assumed that infection confers lifelong immunity against the virus^2,9,15^, but the resulting conclusions about potential outbreak duration may need to be revised if waning of long-term ZIKV antibodies leads to a decline in herd immunity over time. In particular, more work is needed to establish the relationship between waning antibody levels and susceptibility to reinfection, and to be vigilant for potential reemergence of ZIKV in areas that previously had high post-outbreak seroprevalence.

## CONTRIBUTORS

AJK and V-MC-L conceived and designed the study. ADH, MA, MK, AT, CHW, CLL, AJK conducted blood and data collections from the participants. JV and J-CM developed the assays for the serological analyses. AT, TM-H, and TP performed the serological analyses. MA, AT, TM-H, TP, and V-MC-L interpreted the serological data. ADH and AJK performed the statistical analyses. ADH, MA, WJE, JW, V-MC-L and AJK interpreted the statistical results. ADH, MA, V-MC-L and AJK wrote the first version of the manuscript. All authors critically reviewed and approved the final version of the manuscript.

## ACKNOWLEDGMENTS

We are grateful to the minister for education of French Polynesia and to the directors, teachers, nurses and schoolchildren from the elementary and junior high schools selected for the serosurvey in 2018.

We greatly thank all the participants and community leaders in Fiji who generously contributed to the study over the three visits. We would like to acknowledge the work of the field teams: Jessica Paka, Amele Ratevono, Warren Fong, Manisha Prakash, Jonetani Bola, Mosese Ligani, and Taina Naivalu (2017); Meredani Taufa, Adi Kuini Kadi, Jokaveti Vubaya, Colin Michel, Mereani Koroi, Atu Vesikula, and Josateki Raibevu (2015); Dr. Kitione Rawalai, Jeremaia Coriakula, Ilai Koro, Sala Ratulevu, Ala Salesi, Meredani Taufa, and Leone Vunileba (2013), and supported by Eric J Nilles of the World Health Organization Western Pacific Region.

This work was part of R-ZERO Pacific programs funded by the French ministry for Europe and Foreign Affairs [Pacific Funds N°04917-19/07/17]. The study also received support from the French Government’s “Investissement d’Avenir” Program [Labex IBEID N°ANR-10-LABX-62-IBEID], and the Wellcome Trust [Grant number 107778/Z/15/Z]. Fiji 2013 fieldwork was funded by the World Health Organization Western Pacific Region and by the Chadwick Trust. CHW was supported by the UK Medical Research Council (MR/J003999/1). CLL was supported by an Australian National Health and Medical Research Council (NHMRC) Fellowship (1109035). ADH was supported by an MRC LID studentship (grant Number MR/N013638/1). AJK was supported by a Wellcome Trust/Royal Society Sir Henry Dale Fellowship (grant Number 206250/Z/17/Z).

## SUPPLEMENTARY FIGURES AND TABLES

**Supp. Figure 1:**
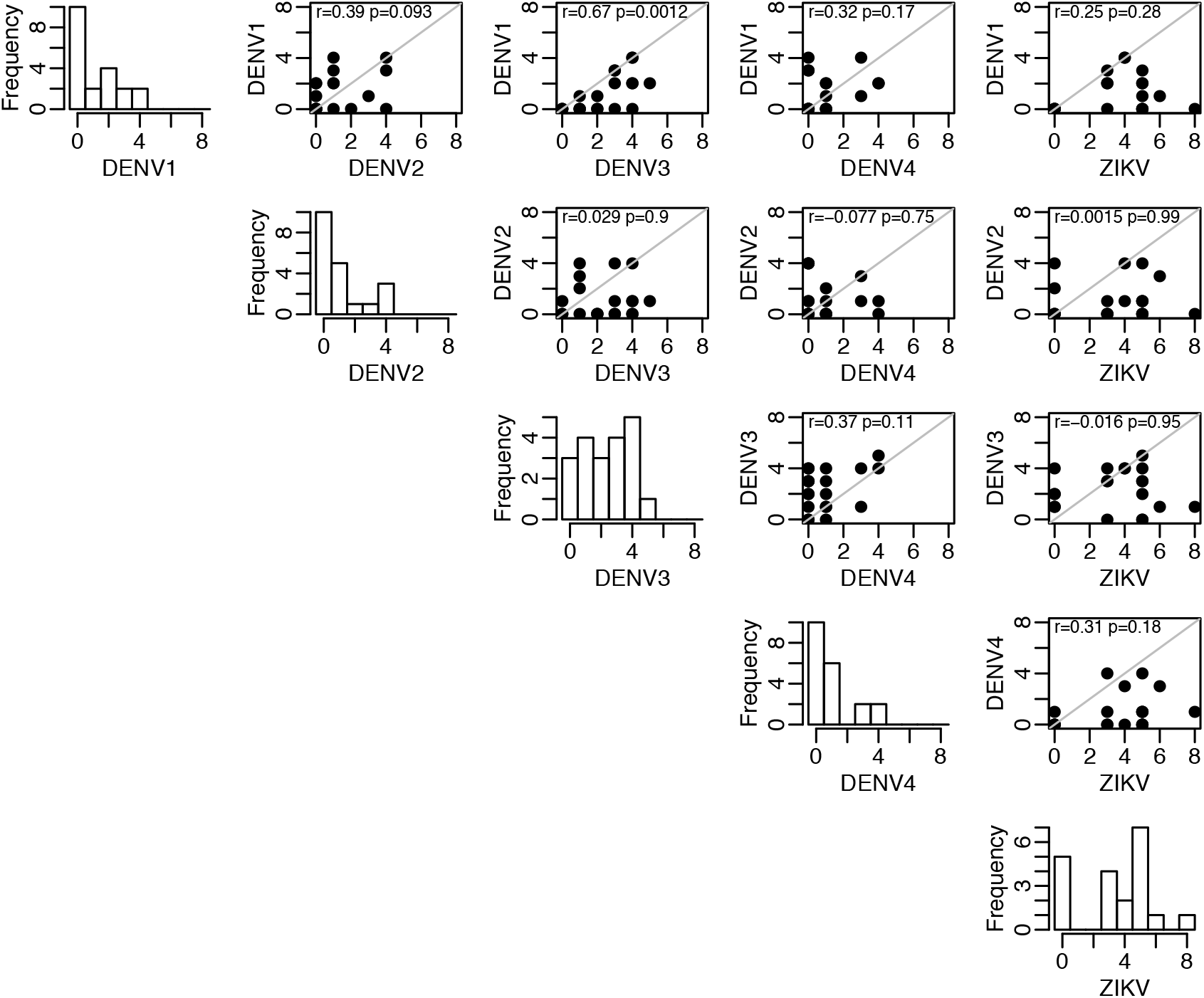
Correlation between rise in DENV and ZIKV log titres between 2013-2015 for individuals who were initially seronegative to all five viruses in 2013. There is significant correlation between DENV-1 and DENV-3 viruses (top row, p=0.0012), which are known to be antigenically similar^1^, suggesting likely crossreactive responses. However, changes in ZIKV titres were not associated with responses to any of DENV viruses (far right column), which strongly indicates that the ZIKV results were genuine infections.

**Supp. Figure 2:**
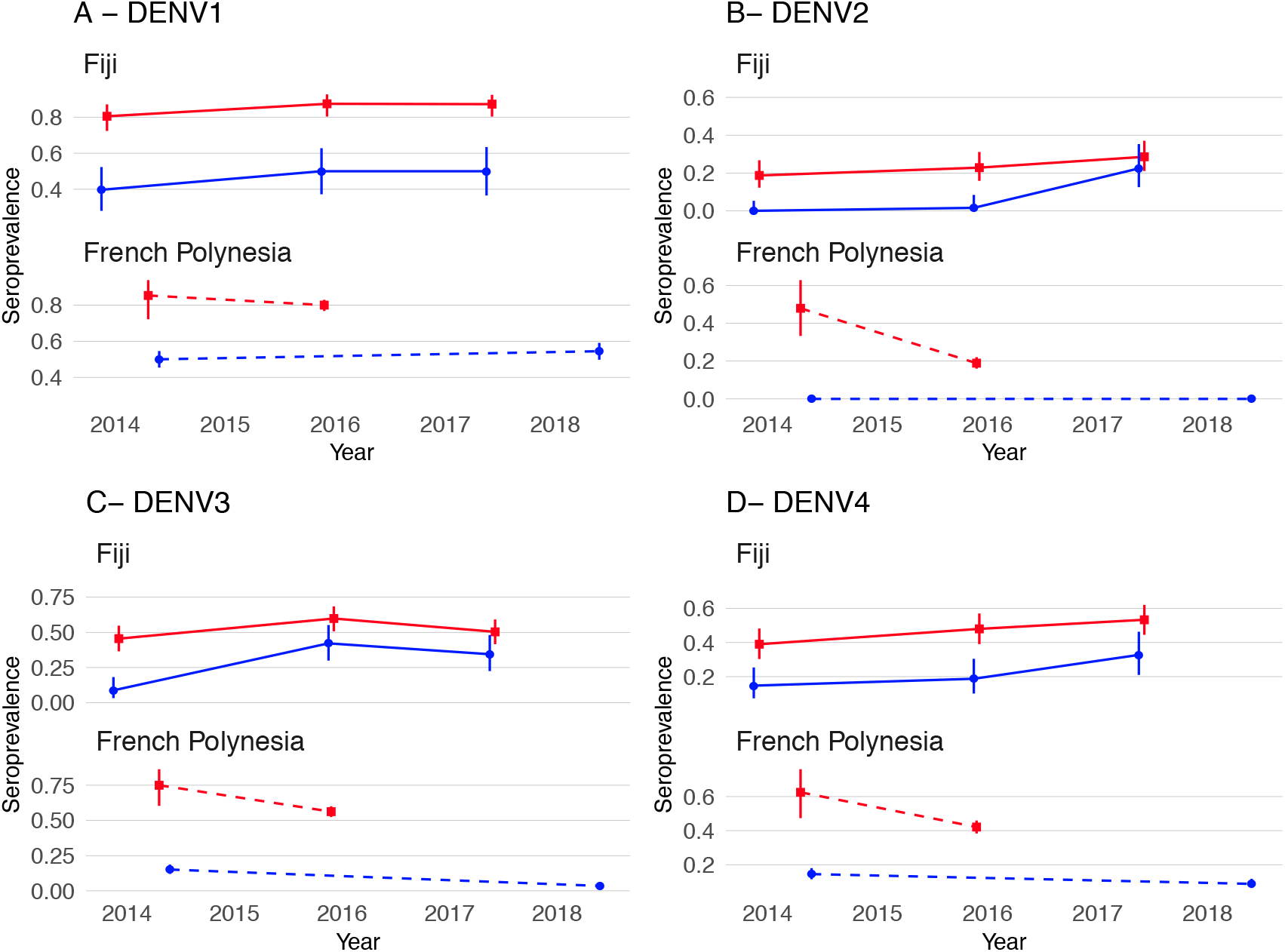
Seroprevalence against each of the four DENV serotypes in Fiji and French Polynesia, by age group. A-D) seroprevalence against DENV serotypes 1-4 in Fiji and French Polynesia, by age group. Red, adults (aged 17+ years). Blue, children (0-16 years). Solid lines, trends in data collected from the same individuals. Dashed lines, trends in seroprevalence between population representative crosssectional surveys. In Fiji, seroprevalence increases between 2015-17 for DENV-1 (A), DENV-2 (B) and DENV-4 (D), and increases for DENV-3 between 2013-15 after a major DENV-3 outbreak^21^ (C). In French Polynesia, between the sampling periods, there were no reported DENV outbreaks for serotypes 2,3,4, and there was hyperendemic DENV-1 circulation. Seroprevalence declines against all four DENV serotypes in French Polynesia (A-D).

**Supp. Figure 3:**
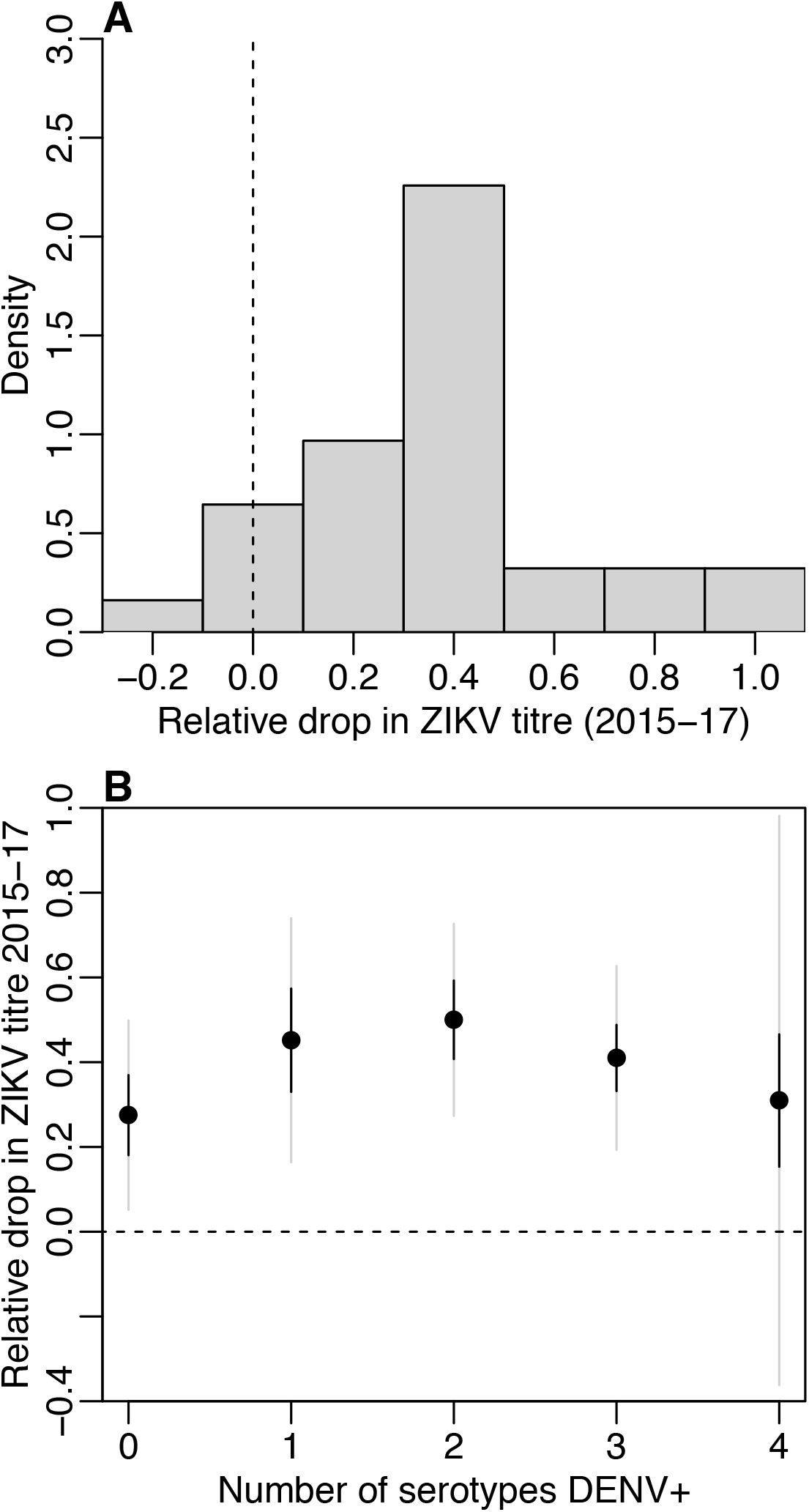
Waning of neutralizing antibody responses against ZIKV, by DENV exposure history. A) Histogram of relative decrease in ZIKV log titre against ZIKV between 2015–2017 for individuals who seroconverted to ZIKV between 2013-2015. B) Relative decline in ZIKV log titre between 2015–2017, stratified by number of prior DENV infections (as measured by seropositivity against that serotype in 2013). Black lines show t-distributed standard error, grey lines show 95% CI.

**Supp. Table 1:**
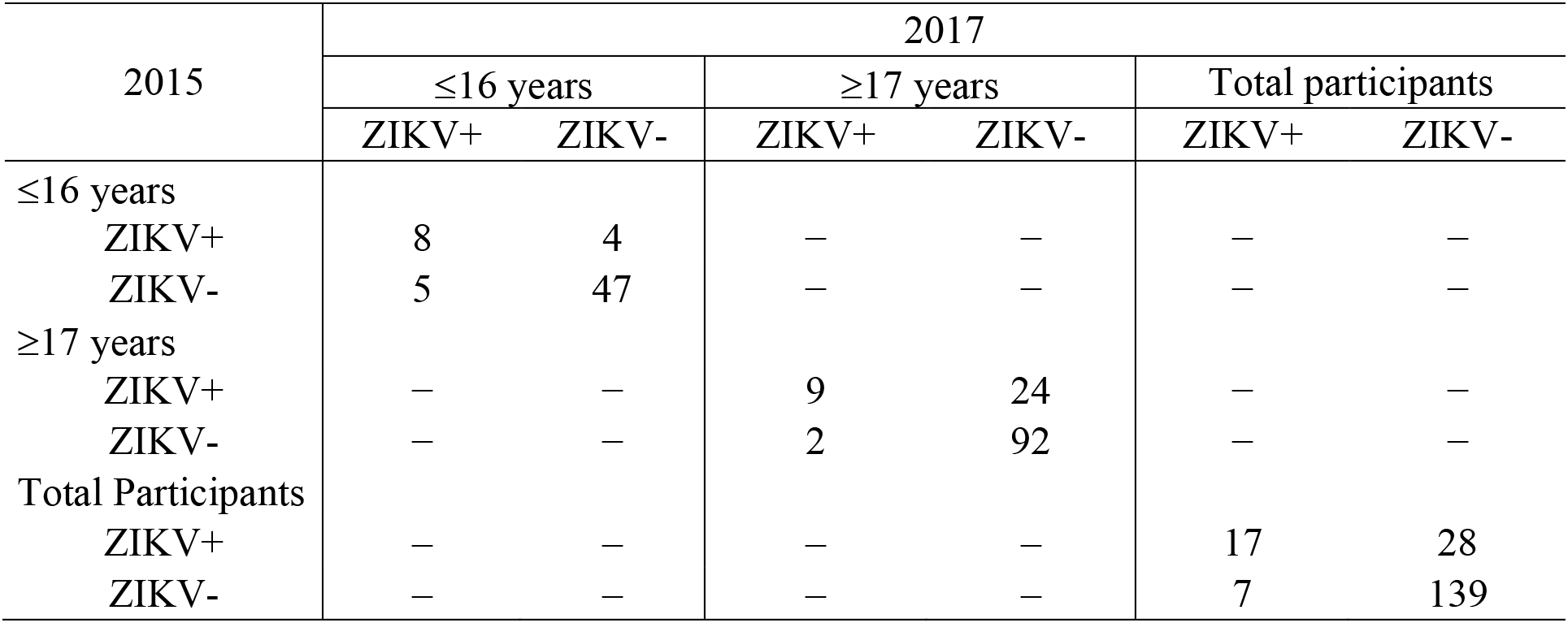
Detection of IgG against ZIKV in the paired samples from participants aged under and over 16 years recruited during October-November 2015 and June 2017 in the Central division in Fiji. Age groups are defined using age of participants when recruited in 2015.

**Supp. Table 2:** Age distribution and profile of DENV exposure history in two crosssectional surveys conducted in the general population from the Society Islands, French Polynesia, in 2014 and 2015. While the age distribution is similar in both studies, the sample in 2014 has a higher proportion of individuals who have tested positive for infection from all four DENV serotypes

| Variable | 2014 (n=49) | 2015 (n=700) |
| --- | --- | --- |
| Age distribution (*median [IQR]*) | 47 [29-56] | 43 [29-57] |
| Number of DENV serotypes positive at time of sample collection (*n [%]*) |  |  |
| 0 | 3 [0·061] | 118 [0·17] |
| 1 | 6 [0·12] | 163 [0·23] |
| 2 | 11 [0·22] | 159 [0·23] |
| 3 | 11 [0·22] | 154 [0·22] |
| 4 | 18 [0·37] | 106 [0·15] |

**Supp. Table 3:** Bootstrap estimated seroprevalence for each of the four DENV serotypes and ZIKV adjusted for sampling bias in two cross-sectional surveys conducted in the general population from the Society Islands, French Polynesia, in 2014 and 2015. Results from the cross-sectional surveys in the Society Islands, French Polynesia, in 2014 and 2015 show a decline in seroprevalence against all 4 DENV serotypes and ZIKV. However, the 2014 sample included more individuals that tested positive for >1 DENV serotype and are assumed to be a higher risk group. We used a bootstrap method with 10,000 iterations which estimated seroprevalence from a sample of the 2014 dataset, taken with replacement, weighted by the DENV exposure distribution in the 2015 survey. After adjusting for the sample bias, there is no evidence of a change in seroprevalence for any of the DENV serotypes but there remains strong evidence that ZIKV seroprevalence declined between 2014-15.

| Virus | 2014<br>seroprevalence<br>(95% CI) | 2014 bootstrap<br>estimates of<br>seroprevalence<br>(95% CI) | 2015<br>seroprevalence<br>(95% CI) | p-value* |
| --- | --- | --- | --- | --- |
| DENV1 | 86 (73-94) | 70 (55-82) | 80 (77-83) | 0·11 |
| DENV2 | 47 (33-62) | 27 (16-39) | 18 (15-21) | 0·2 |
| DENV3 | 76 (61-87) | 55 (41-69) | 55 (51-59) | 1 |
| DENV4 | 63 (48-77) | 43 (29-57) | 42 (38-46) | 1 |
| ZIKV | 37 (23-52) | 42 (29-55) | 22 (19-25) | 0·0047 |

\*chi-squared test comparing 2014 bootstrap estimates with 2015 results

**Supp. Table 4:** Change in neutralization titre between 2013-2017 in a cohort of 46 study participants in Fiji. ZIKV and DENV-3 both circulated between the collection of samples in 2013 and 2015 with ZIKV first reported in July 2015 and DENV-3 circulating between October 2013 and January 2015. Neutralization titre levels rose significantly over this period. Between 2015 and 2017, DENV-3 titre levels still increased with a mean change in tire of 0·89. By contrast, the mean change in ZIKV titre over this period decreased (−1·9).

| Virus | 2013-2015 change,<br>Mean [95% CI] | *p*-value* | 2015-2017 change,<br>Mean [95% CI] | *p*-value* |
| --- | --- | --- | --- | --- |
| ZIKV | 5 [4.5-5.5] | <0.0001 | -1.9 [-2.4-1.5] | <0.0001 |
| DENV3 | 3.4 [2.9-3.9] |  | 0.89 [0.046-1.7] |  |

\* t-test comparing change in neutralization titre for ZIKV and DENV-3 between 2013-2015, and 2015-2017

## Footnotes

1 Katzelnick, L. C., Gresh, L., Halloran, M. E., Mercado, J. C., Kuan, G., Gordon, A.,… Harris, E. (2017). Antibody-dependent enhancement of severe dengue disease in humans. *Science*

